# A new actin depolymerase: a Myosin 1 motor

**DOI:** 10.1101/375923

**Authors:** Julien Pernier, Remy Kusters, Hugo Bousquet, Thibaut Lagny, Antoine Morchain, Jean-François Joanny, Patricia Bassereau, Evelyne Coudrier

## Abstract

The regulation of actin dynamics is essential for various cellular processes. Former evidence suggests a correlation between the function of non-conventional myosin motors and actin dynamics. We investigate the contribution of myosin1b to actin dynamics using sliding motility assays. We observe that sliding on myosin1b immobilized or bound to a fluid bilayer enhances actin depolymerization at the barbed end, while sliding on myosin II, although 5 times faster, has no effect. This work reveals a non-conventional myosin motor as a new type of depolymerase and points to its singular interactions with the actin barbed end.

## INTRODUCTION

Actin filaments (F-actin) form a variety of dynamical architectures that govern cell morphology and cell movements. The dynamics of the actin networks are regulated in space and time by the assembly and disassembly of actin polymers under the control of regulatory proteins. Cortical actin organizes lateral movement of transmembrane proteins and participates in membrane signaling by interacting transiently with the plasma membrane ^1^. One class of actin-associated molecular motors, the single-headed myosin 1 proteins, bridges cortical actin to the plasma membrane. Polymerization of actin filaments at the plasma membrane generates forces on the membrane as well as on their membrane linkers. Inversely myosin 1 can exert and sustain pN forces on F-actin ^2^.

This important class of myosins contains a motor domain at its N-terminus that binds F-actin in response to ATP hydrolysis, a light chain binding domain (LCBD) that binds calmodulin (in most cases), and a Tail domain at the C-terminus (Fig. 1A) ^3^. The Tail domain encompasses a tail homology domain (TH1) with a pleckstrin homology motif (PH) that binds phosphoinositides (Fig. 1A). Beside the involvement of myosin 1 proteins in a large variety of cellular processes including cell migration and membrane trafficking ^3^, manipulation of myosin 1 expression has revealed a correlation between these myosins and actin network architecture ^4–7^. In particular, under- or overexpression of one of these myosins, myosin 1b (Myo1b), affects the organization of the actin cytoskeleton in the juxtanuclear region of HeLa cells ^4^ and in growth cones of cortical neurons ^6^. In contrast to muscle Myosin II (MyoII), this particular Myo1b is a catch-bound motor (the time Myo1b remains bound to F-actin strongly increases with an applied load), it thus remains attached to the filament for a time that depends on the applied force ^8^. Due to its mechanosensitive behavior, Myo1b could in turn exert a force on actin filaments ^8, 9^ and thus affect their polymerization. However, the role of these motors in actin dynamics remains to be explored. In this paper, we use *in vitro* F-actin gliding assays (Fig. 1B) and total internal reflection fluorescence (TIRF) microscopy to study the effect of full-length Myo1b on actin polymerization dynamics, with the motors either immobilized on a solid substrate (Fig. 1B, III) or bound to a fluid supported bilayer, which mimics cell membranes (Fig. 1B, IV).

**Figure 1:**
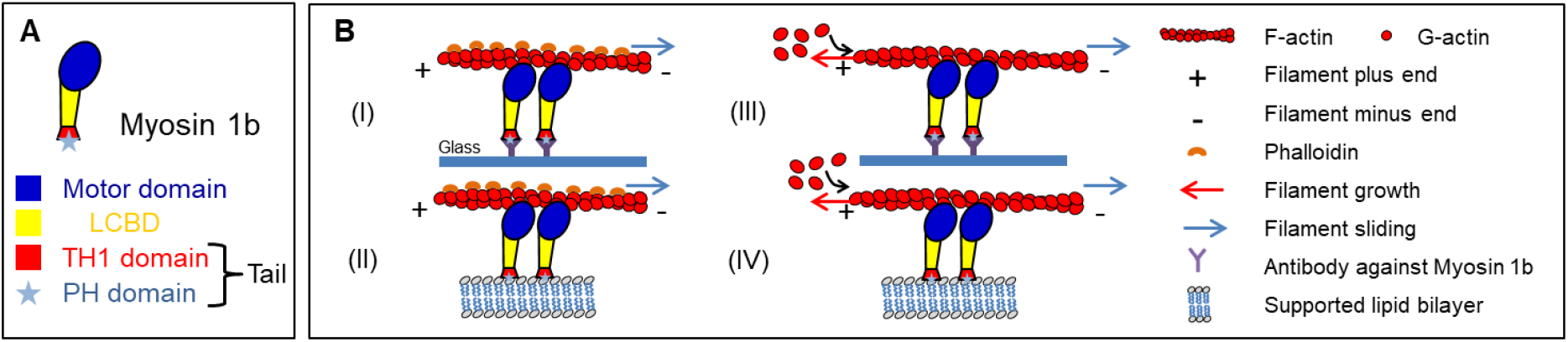
Three-state cross-bridge model and Myo1b-Actin gliding assays. (**A**) Schematic representation of domain organization of Myo1b. Motor domain (blue); Light Chain Binding Domain (LCBD) (yellow); TH1 domain (red), PH domain (cyan) that binds phosphoinositides. (**B**) Gliding assays of stabilized actin filaments (I-II) and polymerizing actin filaments (III-IV) sliding on Myo1b anchored on coverslip (I-III) or bound to a supported lipid bilayer (II-IV).

## RESULTS

We first measured the sliding velocity *v_f_* of single stabilized F-actin on Myo1b immobilized on a glass coverslip (Fig. S1A, top and Movie S1), the sliding velocity *v_f_* and the polymerization rate *v_p_* (expressed in actin sub-unit/s, with the length of an actin subunit being equal to 2.7 nm) of single F-actin (Fig. S1A, bottom and Movie S1) (Materials and Methods), both in the presence of 0.3% methylcellulose for keeping the filaments in the TIRF field, by image analysis. At high Myo1b density (8000 μm^−2^) (for the motor density measurement, see Materials and Methods and Fig. S1B), both stabilized and polymerizing filaments move with the same average sliding velocity *v_f_* = 56.4 ± 15.4 nm.s^−1^ and *v_f_* = 53.9 ± 5.5 nm.s^−1^, respectively (Fig. 2A, Fig.2B, Movie S1 and Table S1) in the presence of 2 mM ATP (above saturation for motor activity) ^10^. In both cases, this velocity decreases by about a factor two when decreasing the Myo1b density by a factor of twenty (Fig. S2B, S2C, Table S1) or when reducing the ATP level to 0.2 mM (Fig. 2A,B, Movies S2, S3) below saturation for Myo1b, but not affecting actin polymerization (Table S2).

**Figure 2:**
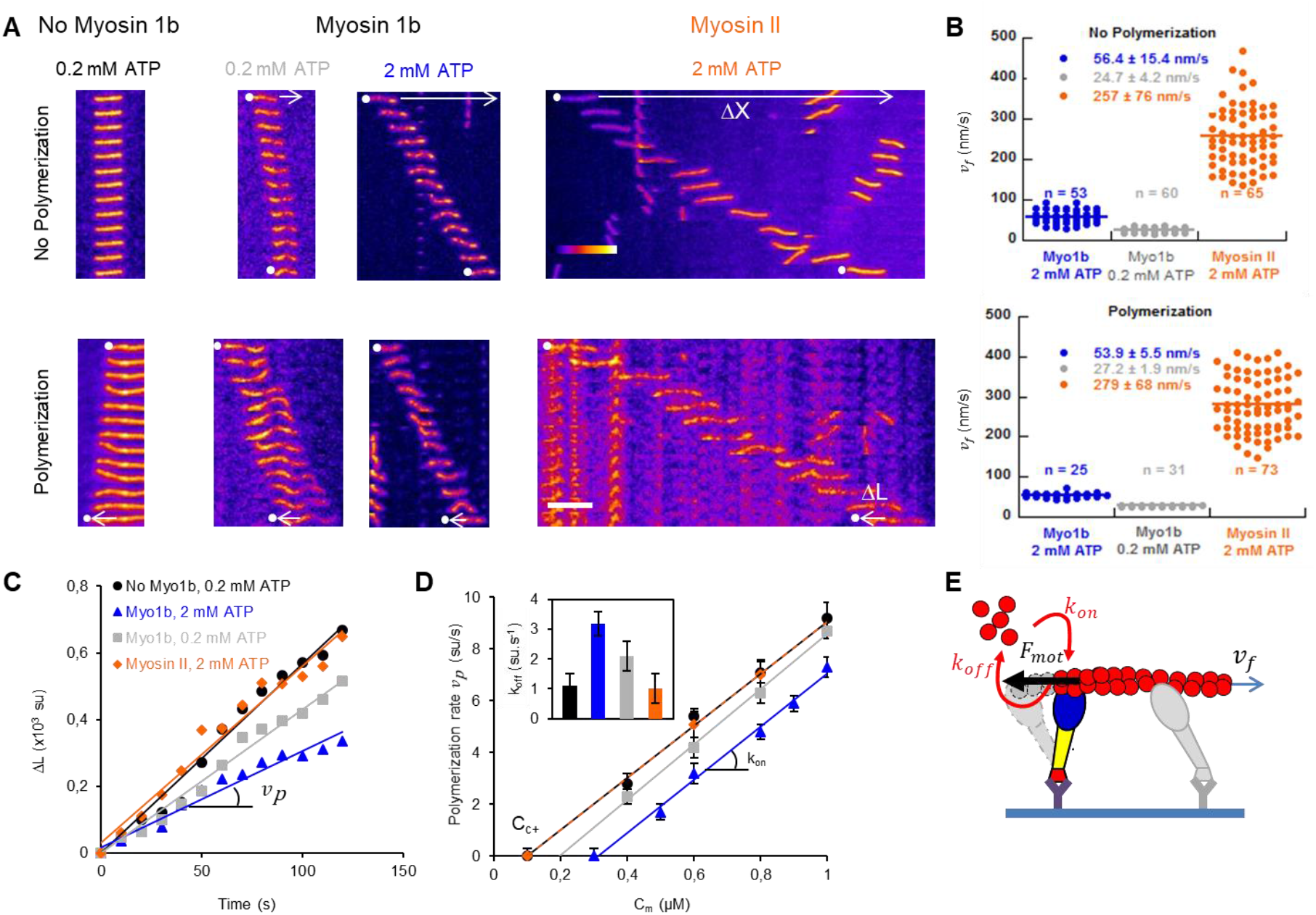
Sliding on immobilized Myosin 1b increases F-actin depolymerization. (**A**) Representative kymographs of stabilized F-actin (top) or polymerizing F-actin with 0.6 μM G-actin (bottom), on uncoated glass or sliding on glass coated with Myo1b (2 mM and 0.2 mM ATP (see movies S2 and S3) or MyoII (see movie S6). The sliding distance *ΔX* and the elongation *ΔL* of the filaments are indicated by white arrows. Actin fluorescence intensity is represented according to the “Fire” LUT of Image J. Scale bar, 5μm. 1 image/10 sec. (**B**) Dot plot representation of the sliding velocities *v_f_* of stabilized (top) and polymerizing actin filaments (0.6 μM G-actin) (bottom) on immobilized Myo1b (8000 molecules/μm^2^) at 2 mM (blue) or 0.2 mM (grey) ATP or sliding on MyoII at 2 mM ATP (orange). The number of analyzed filaments and the mean-values ± s.e.m. are indicated. (**C**) Filament elongation *ΔL* (normalized by the length of the actin subunit (su) equal to 2.7 nm) versus time for filaments shown in A (bottom) in the absence of myosins and in the presence of MyoII or Myo1b at two ATP concentrations. The polymerization rate at the barbed end *v_p_* (in su/s) is deduced from the slope. (D) *v_p_* as a function of G-actin concentration *C_m_* for the different conditions. The fits correspond to *v_p_* = *k_on_C_m_* – *k_off_*, with *k_on_* the rate of association of G-actin and *k_of_* the rate of dissociation. *C*_*c*+_ is the critical concentration for polymerization. Inset: *k_off_* for the different conditions. Error bars represent s.e.m. (n>25). (**E**) Model for the role of Myo1b motor on the dissociation (depolymerization) rate *k_off_*. The filament, sliding at velocity *v_f_*, experiences a force *F_mot_* at the barbed end while the motor is attached, thus impacting *k_off_*, but not the association (polymerization) rate *k_on_*.

We next investigated the impact of Myo1b on actin polymerization upon filament sliding. The actin assembly-disassembly kinetics are an order of magnitude faster at the barbed (plus) end than at the pointed (minus) end ^11^. Thus, we measured the elongation Δ*L* of F-actin at the barbed-end versus time (Fig. 2C). Strikingly, filament sliding on Myo1b decreases the actin polymerization rate *v_p_*, as compared to actin polymerization in the absence of Myo1b (Fig. 2D and Movie S3). This effect is stronger for high filament sliding velocity (in the presence of 2 mM ATP) and weaker at lower Myo1b density on the substrate (Figs. S2B, S2D, Movie S3 and Table S2). We also measured the dynamics of the pointed (minus) end by detecting the relative movement of this extremity compared to a fiducial point on the filament. In contrast with the barbed end, we did not observe any filament length variation (Fig. S2A and Movie S4), thus filament sliding on the motors reduces the actin polymerization rate at the barbed-end only. As a control, we tested the impact on actin polymerization of free Myo1b present only in the bulk, or immobilized on the surface but inactivated (Figs. S2B,D and Movie S5); we did not observe any effect on polymerization (Fig. S2E). Moreover, although actin filaments slide five-fold faster on non- or weak catch-bond myosins such as muscle myosin II (MyoII) ^12^, at the same bulk monomeric-actin (G-actin) concentration (Fig. 2A,B and Movie 56), the actin polymerization rate remains similar to the control (Fig. 2C, D). These observations demonstrate that an immobilized Myo1b motor with intact activity reduces the actin polymerization rate at the barbed-end up to a factor two (Fig. 2D and Table S2) in contrast to muscle MyoII. One important characteristic of Myo1b compared to MyoII that could be relevant, is that it is a catch-bond motor.

Dynamics at the barbed-end results from a balance between the rate of association of G-ctin *k_on_* and the rate of dissociation *k_off_* (Fig. 2E); steady state is obtained at the critical concentration *C*_*c*+_. Classically, these dynamical parameters are deduced from the measurement of the variation of the polymerization rate *v_p_* with G-actin concentration *C_m_*: *v_p_* = *k_on_C_m_* – *k_off_*. By varying the G-actin bulk concentration from 0.1 to 1 μM in the presence of either 0.2 mM and 2 mM ATP, we observed that the slope corresponding to *k_on_* is unchanged when F-actin slides over Myo1b, whereas *C*_*c*+_ which is the ratio between *k_off_* and *k_on_* increases (Fig. 2D) demonstrating that *k_off_* increases under these conditions (Fig. 2D and Table S2). Still, in the absence of G-actin in the bulk, filaments depolymerize faster when they slide over Myo1b (Fig. S2F, G and Movie S7). Interestingly, the dissociation rate is weakly affected when reducing Myo1b density, similarly to sliding velocity (Fig. S2E and Table S2). In contrast, while sliding on MyoII is much faster, this myosin has no influence on *k_off_* at the barbed-end of the filament (Fig. 2D and Table S2). Together, these observations indicate that the Myo1b is an actin depolymerase.

In cells, Myo1b is bound to the fluid plasma membrane lipid bilayer through the interaction of its PH domain with PI(4,5)P2 ^13^, and thus it is not immobilized. We mimic experimentally these cellular conditions by analyzing the impact of Myo1b on actin dynamics when bound to a glass-supported lipid bilayer (SLB) composed of 79.5% POPC, 20% L-α-phosphatidylinositol-4,5-bisphosphate (PI(4,5)P2) and 0.5% Rhodamine-PE or Atto488-DOPE (mol/mol) (Fig. 1B-IV and Fig. 3) (Materials and Methods). We checked using fluorescence recovery after photobleaching (FRAP) that membrane fluidity was preserved in the SLB with bound Myo1b (Fig. 3A and Fig. S3). The lipid diffusion coefficient was in agreement with data published on SLBs composed of pure POPC ^14^. After recruitment on the SLB, Myo1b diffuses freely in the plane of the membrane (Fig. 3A). We did not observe any difference between experiments with or without methylcellulose in the bulk (Fig. 3A). In addition, the lipids continue to diffuse freely even when Myo1b diffusion is strongly decreased by a dense actin network (Fig. 3A) due to an emerging coupling when a filament bridges multiple motors. The diffusion coefficients are close to those measured in cell membranes (Fig. 3A), showing that in our *in vitro* experiments, the fluidity of the membrane is preserved. As previously reported ^15^, myosin 1 proteins bound to a lipid bilayer exert a force strong enough to propel actin filaments in spite of the fluidity of the support. We confirmed that in the presence of 2 mM ATP and at a similar Myo1b density as when immobilized (8500 μm^−2^), stabilized and polymerizing F-actin slides on Myo1b bound to SLBs, although with a velocity reduced by about 25%: *v_f_* = 37.6 ± 7.3nm.s^−1^ and *v_f_* = 39.3 ± 8.2nm.s^−1^ respectively (Fig. 3B, Fig. 3C, Movie S8 and Table S1).

**Figure 3:**
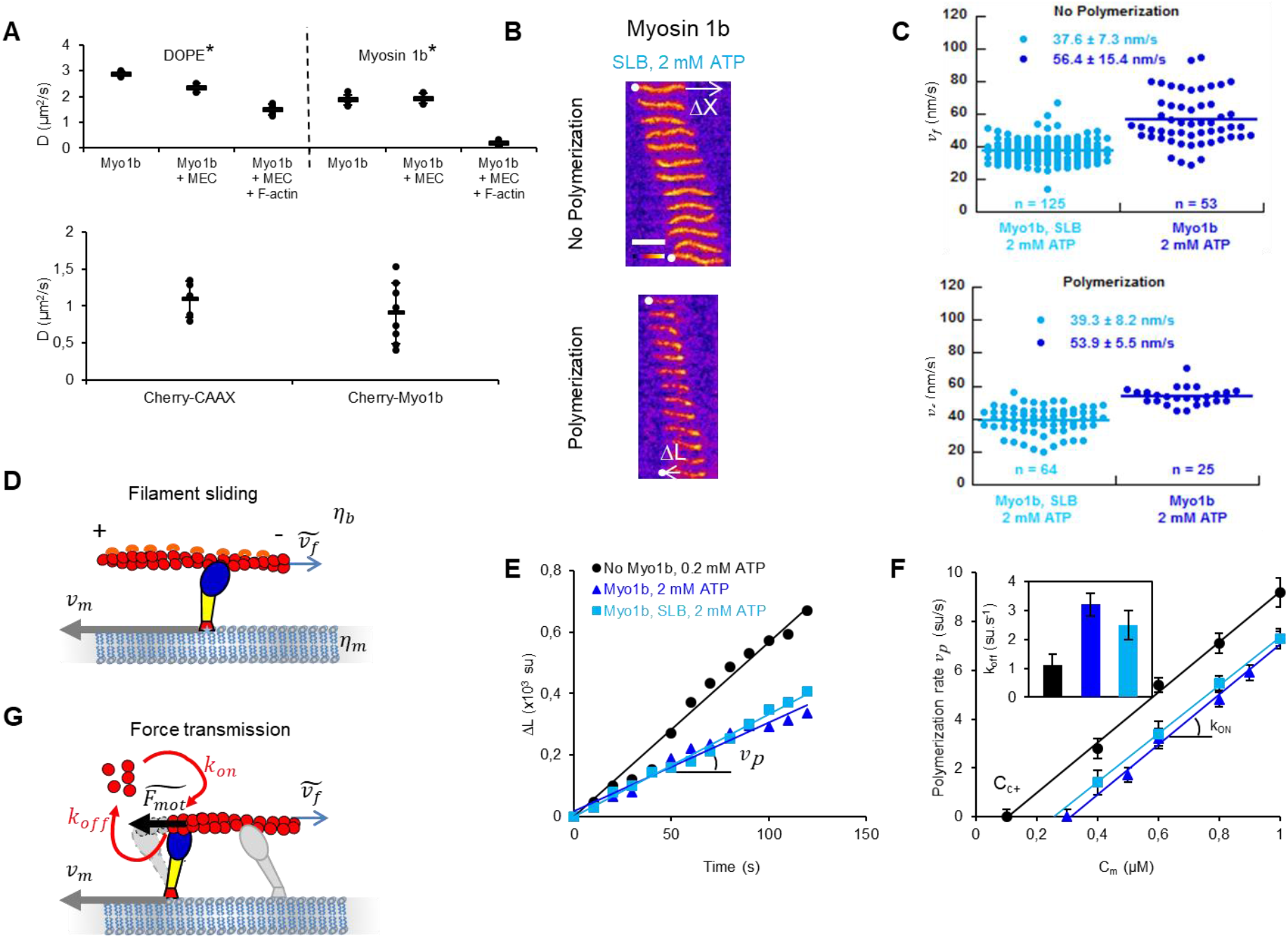
Sliding on Myosin 1b bound to a supported lipid bilayer increases F-actin depolymerization. **(A)** Top: Diffusion coefficients of Atto488DOPE (DOPE*) and Alexa488-labelled Myo1b (Myo1b*) in a SLB with bound Myo1b, with or without 0.3 % methylcellulose (MEC), and in absence or in the presence of a dense F-actin network (n = 30). See Fig. S7 for representative FRAP experiments. Bottom: Effective diffusion coefficients of Cherry-CAAX, Cherry-Myo1b, expressed in HEK293T cells (n > 5). Error bars represent s.e.m. **(B)** Representative kymographs of non-polymerizing (top) and polymerizing F-actin (bottom) in the presence of 0.6 μM G-actin with Myo1b bound to SLBs (movie S8). Scale bar, 5μm. 1 image/10 sec. **(C)** Dot blot representation of the velocities *v_f_* of stabilized (top) and polymerizing F-actin (bottom) sliding on immobilized Myo1b (dark blue) or on Myo1b bound to a SLB (cyan). The number of analyzed filaments is indicated. **D)** Model for filament sliding: The effective filament sliding is determined by a balance between the viscous dissipation of the motor moving with a velocity *v_m_* in the lipid bilayer with a viscosity *η_m_* and a filament sliding at a velocity 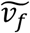 in a solution of viscosity *η_b_*. **(E)** Δ*L* versus time for the single filaments shown in (B). **(F)** *v_p_* as a function of G-actin concentration *C_m_* for the different conditions. The fit to the data is the same as in Fig. 2D. Inset: *k_off_* for the different conditions. Error bars represent s.e.m. (n>25). **(G)** Model for force transmission: The effective force experienced by the polymerizing filament 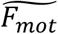 is diminished by the motion in the lipid bilayer of the motor *v_m_*at the barbed end.

We have calculated the relative contributions of the viscous drag of the bulk and of the lipid bilayer on the motion of the filaments (Supplementary Inf.). First, we have considered F-actin moving in water (*η_b_* = 10^−3^*Pa. s*) above Myo1b bound to a SLB (Fig. 3D). We estimate that, since the in-plane viscous drag between the motor and the lipid bilayer is much larger than the bulk viscosity experienced by the actin filaments, the velocity of the motors bound to actin filaments with respect to the bilayer couple, *v_m_*, practically vanishes. Thus, filaments slide with a velocity 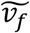 similar to that measured for immobilized motors: 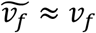 (Fig. S4). Including the increased viscosity of the bulk in the presence of methylcellulose (10^−2^ Pa.s at 0.3%, product information Sigma) and crowding effects between nearby filaments reduces the effective sliding speed of the filament 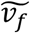 since part of the sliding is dissipated by in-plane motion of the motors in the bilayer (Fig. S4). This can explain why in our experiments, F-actin moves over SLB-bound Myo1b but with a slightly reduced velocity as compared to immobilized Myo1b (Fig. 3C, Table S1). This is in line with the results by Grover et al ^16^ showing a decreased gliding velocity of membrane-anchored kinesins due to their slippage in the lipid bilayer.

In these experimental conditions, we observed a significant increase of the actin depolymerization rate at the barbed end *k_off_* when filaments slide on Myo1b bound to a SLB, although weaker than for immobilized Myo1b, while keeping the polymerization rate unchanged (Fig. 3E, Fig. 3F and Table S2). We conclude that the dissipation of sliding filaments in SLBs is low enough to let Myo1b exert a significant dissociation force even when bound to a fluid membrane (See force balance in Fig. 3G).

## DISCUSSION

As previously shown, MyoII induces actin network contraction, potentially leading to filament buckling and breaking ^17, 18^. However, we show here that muscle MyoII does not affect actin polymerization dynamics. Different actin-binding proteins are already known for preventing actin polymerization (capping protein) ^11^, enhancing it (formin) ^19, 20^ or depolymerizing actin (ADF/cofilin) ^21, 22^ at the barbed end. Also, some kinesin motors, e.g., kinesins 8 and 13, have been shown to depolymerize microtubules ^23, 24^. We demonstrate here that Myo1b, but not MyoII, induces a significant actin depolymerization at the barbed end (Tables S1 and S2). This suggests that a mechanical process is involved in actin monomers’ removal at the actin filament tip. One possible mechanism could be through the modulation of the torsion of the filaments ^25^. In this case, the polymerization kinetics is expected to depend on the filament length with a twist gradient inversely proportional to the length. However, this is not what we observe (Fig. S2H), excluding an explicit role of filament torsion due to motor attachment along the filament. Since the effect is essentially detected at the extremity of the filament, a local process should account for depolymerization by Myo1b. Given the motor activity of Myo1b, the increased actin depolymerization could be of mechanical origin and due to a force exerted on the filament by the last Myo1b motor close to its barbed end. Any mechanism where a motor at the barbed end has a longer attachment time would create a force opposing the motion of the actin filament that could lead to an enhanced actin dissociation at this extremity. This force and therefore the depolymerization rate increase with the filament velocity and with the attachment time of the last Myo1b to the filament. We could for instance consider that Myo1b has specific molecular properties when bound at the barbed end, such as a higher affinity or a different type of interaction with this part of the filament. The stronger binding could also have a mechanical origin, such as a catch-bond effect. Myo1b is indeed a strong catch-bond motor in the few pN force range ^8^, as compared to MyoII ^12^. However, single molecule experiments have evidenced that the motor has an enhanced attachment time when it resists to a load ^8^, but this does not correspond to our experimental situation. Very few experiments have been performed with assisting loads ^26^, only on a very low force range, not with the full length Myo1b and far from actin extremity; thus, some unexpected load-induced detachment reduction might occur in our conditions. Experiments that could provide molecular details on the interaction of Myo1b with actin filaments plus-end are obviously extremely challenging, but nevertheless, we have uncovered a peculiar behavior of Myo1b at this filament tip with our gliding assay.

These observations indicating that Myo1b is an actin depolymerase, even when bound to a lipid bilayer, suggest together that this myosin is able to regulate actin dynamics *in vivo* nearby different cellular membranes. Myo1b’s influence on actin dynamics can control the organization of actin networks, as reported in growth cones ^6^. An actin network can be impacted by Myo1b in different ways. It can reduce the length of actin filaments, as shown by this work, and thus change the mesh-size, or the cortical thickness and consequently the cortical contractibility ^27^. Whether or not it can affect the Arp2/3-dependent branched actin network and/or formin-dependent actin bundles remains to be explored. Moreover, since Myo1b is specifically present at the interface between the plasma membrane and the cortical actin, Myo1b may coordinate receptor signaling by arranging the cytoskeleton ^28^. Further experiments need to be performed in the future to determine the relative contribution of Myo1b with respect to the other proteins that regulate actin dynamics.

Experimental evidence supports a role of several Myosin 1 proteins in membrane remodeling ^3^. Similarly to capping proteins ^29^, Myo1b and perhaps other Myosin 1 proteins could shape membranes by regulating the growth of filaments at the plasma membrane. Alternatively, Myo1b could shape membranes by inducing stresses in the cortical actin. Indeed, Myo1b induces actin movement and reduces actin growth when bound to supported bilayers, as shown in our experiments. Since the fluidity of our synthetic membranes and of cellular membranes are similar (Fig. 3A), we propose that Myo1b has the same function in cells. Collectively, these motors could drive the sliding of actin filaments at the membrane surface, which could create stresses that relax by deforming the cortex and the attached membrane. Interestingly, when Myo1b is bound to a deformable giant liposome, we observed that it produces membrane invaginations in presence of stabilized actin filaments (Fig. S5).

Besides myosin II and myosin 1 proteins, myosin VI has also been reported to influence the actin architecture during, e.g. spermatid individualization in Drosophila ^30^ or around melanosomes ^31^. It might be time now to take a fresh look on the involvement of non-conventional myosins in actin dynamics and organization.

## Materials and Methods

### Protein purification

Actin was purified from rabbit muscle and isolated in monomeric form in G buffer (5 mM Tris-HCl, pH 7.8, 0.1 mM CaCl_2_, 0.2 mM ATP, 1 mM DTT and 0.01% NaN_3_). Actin was labeled with Alexa 594 succimidyl ester-NHS ^32^

Myosin II was purified from rabbit muscle as previously described ^33^.

Expression and purification of Myosin 1b: FLAG-myo1b was expressed in HEK293-Flp-In cells cultured in Dulbecco’s modified Eagle medium supplemented with 10% fetal bovine serum and 0.18 mg ml^−1^ hygromycine in a spinner flask at 37 °C under 5% CO_2_, and collected by centrifugation (1,000 g, 10min, 4 °C) to obtain a 4–5 g of cell pellet. The pellet was lysed in FLAG Trap binding buffer (30 mM HEPES, pH 7.5, 100 mM KCl, 1 mM MgCl_2_ 1mM EGTA, 1 mM ATP, 1 mM DTT, 0.1% protease inhibitor cocktail (PIC), 1% Triton X-100) for 30 min at 4 °C and centrifuged at 3,400 g for 10 min at 4 °C. The collected supernatant was then ultracentrifuged (250,000 g, 60 min, 4 °C). The solution between pellet and floating lipid layer was incubated with 150 μl of anti-FLAG beads for 2 h at 4 °C. The beads were collected by centrifugation (1,000 g, 5 min, 4 °C). After a washing step, FLAG-myo1b was then eluted by incubating with 0.24 mg ml^−1^ of 3X FLAG peptide in 300 μl elution buffer (binding buffer without Triton X-100 supplemented with 0.1% methylcellulose) for 3 h at 4 °C. After removal of the beads by centrifugation (1,000 g, 3 min, 4 °C), the protein solution was dialyzed against elution buffer overnight at 4 °C to remove the 3X FLAG peptide. Myo1b was fluorescently labeled using Alexa Fluor 488 5-SDP ester ^34^. Inactivated Myo1b was removed by ultracentrifugation (90,000 rpm, 20 min, 4 °C) with 10 μM F-actin in presence of 2 mM ATP. Inactivated Myo1b was then dissociated from F-actin by incubating the pellet collected after untracentrifugation in elution buffer (30 mM HEPES, pH 7.5, 100 mM KCl, 1 mM MgCl_2_, 1mM EGTA, 1 mM ATP, 1 mM DTT and 0.1% methylcellulose) supplemented with 1 M NaCl and collected in the supernatant after a second centrifugation (90,000 rpm, 20 min, 4 °C).

### Supported lipid bilayer (SLB) preparation

SLBs were formed by fusion of small unilamellar vesicles (SUVs) prepared as follows. Lipid mixtures containing 79.5 % POPC, 20 % L-α-phosphatidylinositol-4,5-bisphosphate (PI(4,5)P2) and 0.5 % Rhodamine-PE or Atto488-DOPE (mol/mol) were mixed together in a glass vial, dried with N_2_, placed in vacuum desiccator for 1 hour, then rehydrated with Fluo F buffer (5 mM Tris-HCl-pH 7.8, 100 mM KCl, 1 mM MgCl_2_ 0.2 mM EGTA, 0.2 mM or 2 mM ATP, 10 mM DTT, 1 mM DABCO, 0.01% NaN_3_) for 30 min at room temperature, to a final lipid concentration of 2 mg/mL. After rehydration, the glass vial was vortexed to detach the liposomes. SUVs were formed by sonication, aliquoted and stored at −20 °C. For SLB formation by fusion, CaCl_2_ was added to a final concentration of 5 mM, with 50 μl of SUVs. The solution was incubated in the chamber for 20 min and washed 5 times with Fluo F buffer 0.1 % BSA. The quality of the SLB was checked by FRAP.

### Giant unilamellar vesicle (GUV) preparation

Lipid compositions for GUVs were 79.7 % POPC, 20 % L-α-phosphatidylinositol-4,5-bisphosphate (PI(4,5)P_2_) and 0.3 % Texas Red DHPE. GUVs were prepared by using polyvinyl alcohol (PVA) gel-assisted method in a 200 mM sucrose buffer at room temperature for 2 hour as described previously ^35^.

### Myosin 1b surface density

We measured the protein surface density (number of proteins per unit area) on solid surfaces or on SLBs by using a previously established procedure ^36, 37^. It is calculated from a labeled proteins/lipids calibration. We first measure the fluorescence of POPC SLBs containing predefined amounts of Atto488-DOPE fluorescent lipids (DOPE*) to establish the relationship between the density of DOPE* *n_DOPE*_* and the corresponding fluorescence intensity 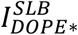 (Fig. S1Ba). Assuming an area per POPC of 0.68 nm^2^, we derive the calibration coefficient A corresponding to the slope of this curve. Note that A depends on the illumination and recording settings of the microscope.

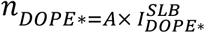

Since Myo1b is labeled with Alexa488 and not Atto488, we have to correct this value by the ratio of fluorescence of the two fluorescent dyes in bulk deduced from the slope of the titration curves 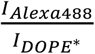 (Fig. S1Bb and c). We then obtained the surface density of the protein deduced from the measurement of the Myo1b-Alexa488 intensity *I*_*Myo*1*b*\*_ as: 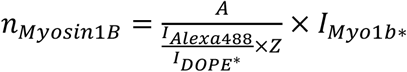 where *Z* is the degree of labeling for the protein of interest (Here, *Z*=1). In our experiments, the calibration factor 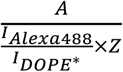 is equal to 0.278.

### Single-filament TIRF microscopy assays

The kinetics of single filament assembly was monitored by TIRF microscopy (Eclipse Ti inverted microscope, 100X TIRF objectives, Quantem 512SC camera). The experiments were controlled using the Metamorph software. Coverslips and glass slides were sequentially cleaned by sonication with H2O, ethanol, acetone for 10 min, then 1M KOH for 20 min and H2O for 10 min. In the case of supported lipid bilayer, first the coverslips and glass slides were cleaned by sonication with Hellmanex III (Hellma Analytics) for 30 min. Flow chambers were assembled with a coverslip bound to a glass slide with two parallel double-stick tapes. The chamber was incubated with 100 nM anti-myo1b antibody in G buffer (5 mM Tris-HCl, pH 7.8, 0.1 mM CaCl_2_, 0.2 mM ATP, 1 mM DTT and 0.01% NaN_3_) for 10 min at room temperature. The chamber was rinsed three times with buffer G 0.1 % BSA and incubated 5 min at room temperature. Then the chamber was incubated with 300 nM Alexa488-labeled myo1b in Fluo F buffer (5 mM Tris-HCl, pH 7.8, 100 mM KCl, 1 mM MgCl_2_ 0.2 mM EGTA, 0.2 mM or 2 mM ATP, 10 mM DTT, 1 mM DABCO, 0.01% NaN_3_) for 10 min at room temperature. Assays were performed in Fluo F buffer, containing 0.2 or 2 mM constant ATP, supplemented with 0.3% methylcellulose (Sigma) and with G-actin (10 % Alexa594) or F-actin (stabilized with phalloidin-Alexa594) at indicated concentrations. To maintaining a constant concentration of ATP in this assay an ATP regenerating mix, including 2 mM ATP, 2 mM MgCl_2_, 10 mM creatine phosphate and 3.5 U/mL creatine phosphokinase, which constantly re-phosphorylates ADP into ATP to maintain a constant concentration of free ATP, was added.

The sliding and elongation velocities of actin filaments were analyzed by using Kymo Tool Box plugin of Image J software (https://github.com/fabricecordelieres/IJ_KymoToolBox). Only filaments longer than 20 pixels are analyzed. When filaments slide on myosins, only those moving directionally during the whole sequence are selected. On each image of a sequence, a segmented line is manually drawn over a single filament, which generates a 10 pixel wide band. The plugin flattens the curved filaments and generates a kymograph. The accuracy on the displacement and the length of the filaments is of the order of the pixel size (160 nm). We consider that each actin subunit contributes to 2.7 nm of the filament length.

### FRAP methods

For diffusion measurements, Fluorescence Recovery After Photobleaching (FRAP) experiments were performed through a X100 or X60 oil immersion objective on an inverted spinning disk confocal microscope (Nikon eclipse Ti-E equipped with a Prime 95B^™^ Scientific CMOS camera, Photometrics) equipped with a FRAP unit. Recovery curves (average of 5 independent experiments, performed on different circular regions of the SLB using the same bleaching conditions) were normalized to the initial intensity and fitted with a single exponential function. We derive the *τ*_1/2_ time corresponding to the time at which the fluorescence signal has recovered 50% of its value before bleach. We calculated the diffusion coefficient using the Soumpasis equation ^38^: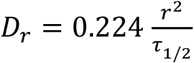, where r is the radius of the bleached region.

## Acknowledgments

We thank B. Goud for insightful discussions, C. Le Clainche (I2BC, Gif-sur-Yvette, France) for providing actin and Myosin II and critically reading the manuscript, F.-C. Tsai for SLB preparation, L. Blanchoin, C. Leduc, J. Prost, M. Henderson for carefully reading the manuscript. The authors greatly acknowledge the Cell and Tissue Imaging (PICT-IBiSA), Institut Curie, member of the French National Research Infrastructure France-BioImaging (ANR10-INBS-04). This work was supported by Institut Curie, Centre National de la Recherche Scientifique (CNRS), the European Research Council (ERC) (J.F.J., P.B. and E.C are partners of the advanced grant, project 339847 and their groups belong to the CNRS consortium CellTiss, the Labex CelTisPhyBio (ANR-11-LABX0038) and Paris Sciences et Lettres (ANR-10-IDEX-0001-02). J.P. and R.K. were funded by the ERC project 339847.

## Author contributions

P.B and E.C. designed the study. J.P. and A.M. performed TIRF experiments and analyzed data; J.P. and T.L. conducted FRAP experiments; H.B. purified Myosin 1b. R.K. and J.-F.J. developed the model. P.B, E.C., J.P., R.K. and J.-F.J. wrote the paper.

## Supplementary Information

### Filament sliding on a lipid bilayer

In order to estimate how the motor force is transmitted to the filament when the molecular motors are immersed in a lipid bilayer, instead of being rigidly anchored to a solid surface, we write a simplied force balance between the viscous friction force of the motor/filament and the force exerted by the molecular motors *nF_mot_* where *n* = *ρl* is the number of attached motors along the filament of length *I*,

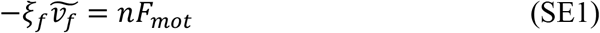

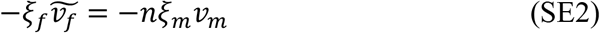

where *ξ_m_* is the in-plane friction coeffcient of the motor complex in the lipid bilayer, *V_m_*, the speed of a molecular motor, *ξ_f_*, the friction coefficient between the filament and the surrounding solution and 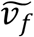, the speed of the filament in the assay. The first equation is the force balance on the filament and the second equation is the force balance on the filament and motors complex. We use here a simplified expression for the motor force ^39^,

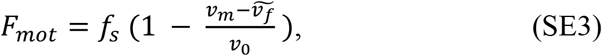

where *f_s_* is the stall force of one motor and *v_0_* the motor speed at vanishing external force (*f_s_* ≈ 0 (1) pN and *v_0_* ≈ 50 *nm/s*). The friction between the filament and the solution can be estimated as *ξ_f_* ≈ 2*Πn_b_*/ (log l/*b_f_*) ≈ *O* (10^−8^)*Pa. s. m*, where we use the bulk viscosity of water *n_b_* = *O* (10^−3^)*Pa.s* and as a cut-off lengthscale, the size of the filament *l* = *O* (10^−5^)*m*. Note however that the effective bulk viscosity can be significantly larger since the filament slides close to a surface. The friction between the motor complex and the lipid membrane is *ξ_m_* ≈ 4*Πn_m_*/ (log l_0_/*L*) ≈ *O* (10^−9^)*Pa. s.m* ^40^, where *L* is the size of the membrane and l_0_ the size of a motor (we estimate the membrane viscosity as *n_m_* ≈ O (10^−10^)*Pa.s* ^40^ and (log l_0_/*L*) = *0* (1)). Solving equations SE 1 and SE 2 gives the following values for the velocity of the filament, 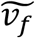, relative to the velocity at zero external force on a solid substrate, *v*_0_,

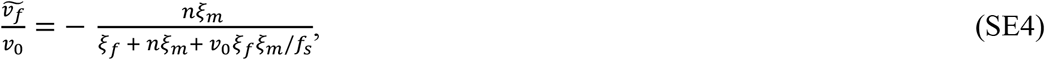

and for the velocity of the motor, *v_m_*,

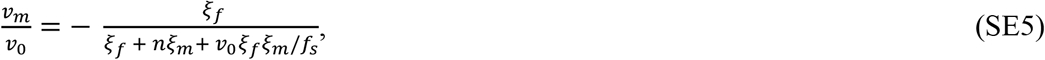

For realistic values of the friction coefficient of water and typical force values we obtain a filament speed which is very close to the filament speed on a solid substrate 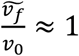, indicating that, since the in-plane membrane friction of the motor is larger than the filament friction with the fluid, the motors are effectively immobile. However, upon increasing the viscous friction between the filament and the bulk by one/two orders of magnitude, potentially due to inter-filament friction (at high filament density) or to the addition of methylcellulose, the sliding speed of the filament diminishes significantly (Fig. S4). Also decreasing the density of motors along the filament impacts the sliding speed since the effective friction between membrane and motor is proportional to the density of motors.

## Legends - Supplementary tables and figures

**Table S1:**
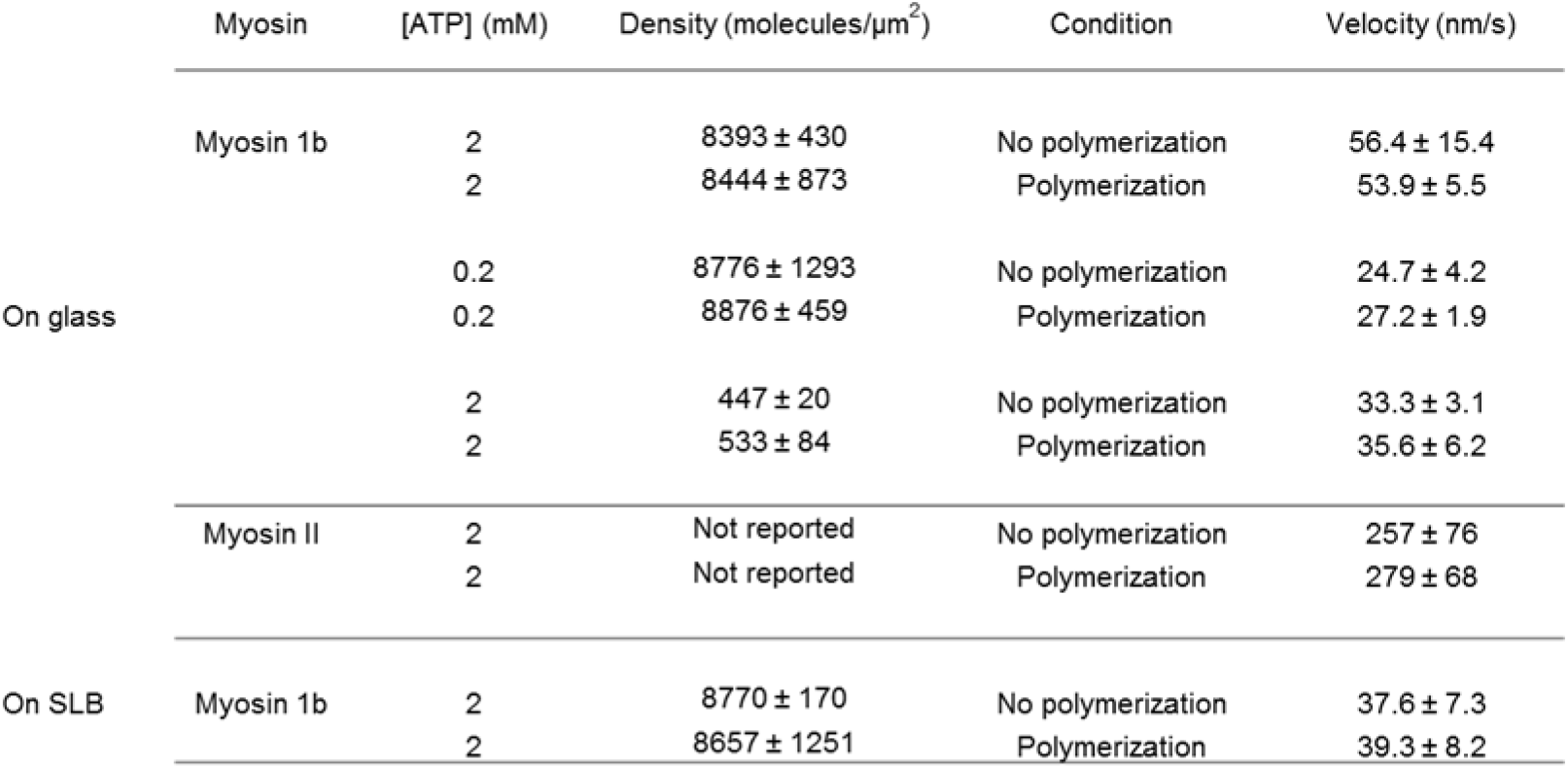
Sliding velocities *v_f_* of stabilized and polymerizing actin filaments on Myo1b or Myosin II in the different used conditions.

**Table S2:**
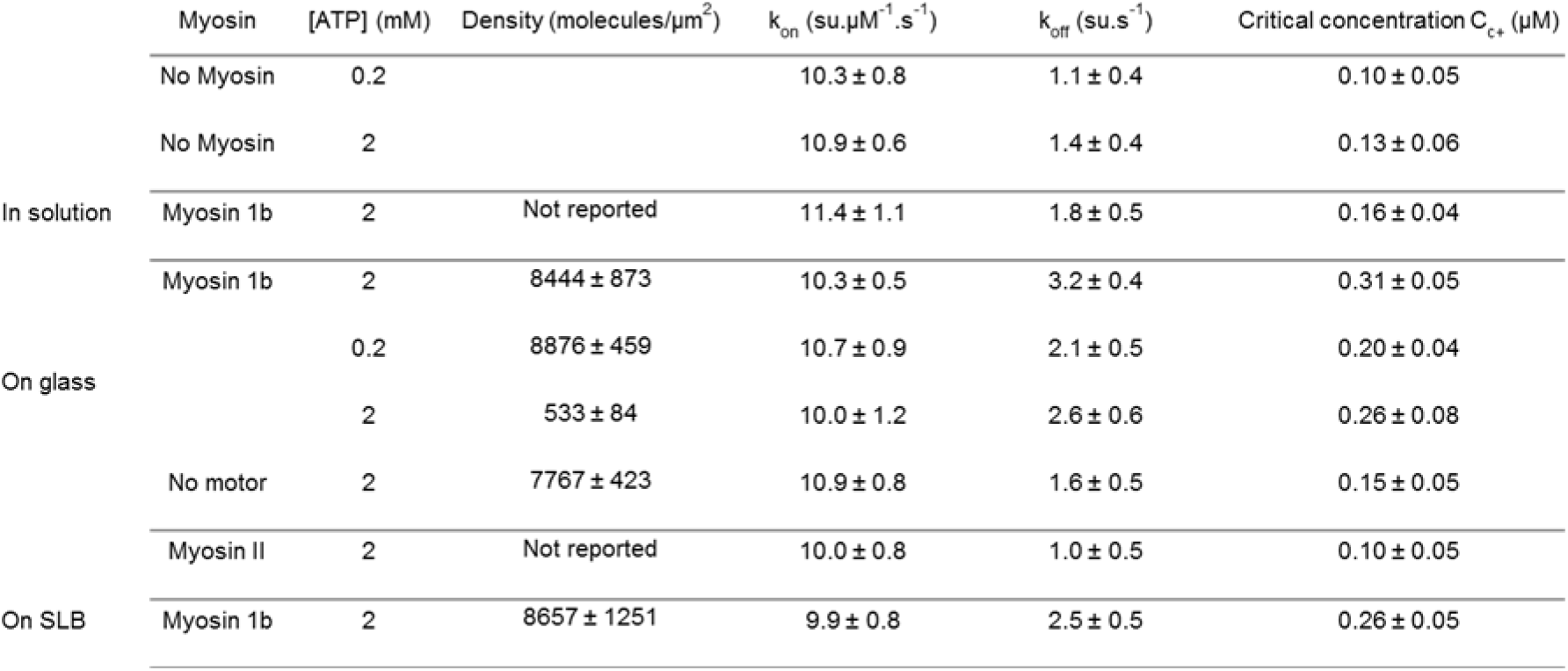
Rate constants of G-actin monomer association and dissociation in the absence and presence of Myo1b or Myosin II in the different used conditions.

**Figure S1:**
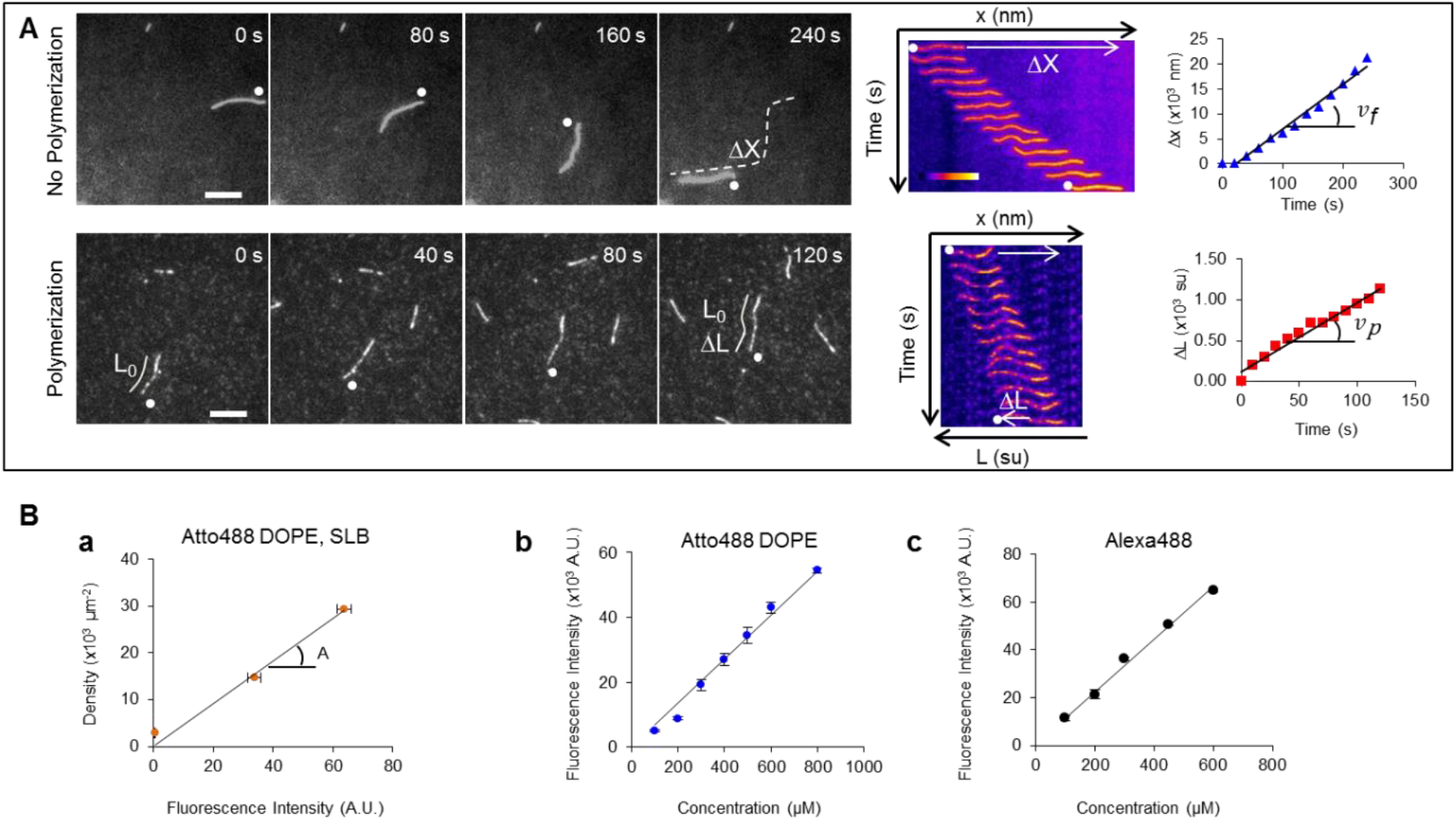
Analysis of the experimental data and calibration of the Myo1b density. **(A)** Left: Time-lapse images obtained by TIRF microscopy of a stabilized filament (top) (and a polymerizing filament in the presence of 1.2 μM G-actin (bottom) sliding along glass-anchored Myo1b (see movie S1). White dots indicate the filament’s barbed end. The white dashed represents the trajectory of the stabilized filament, and *ΔX* the total displacement of the filament over the period considered. *L_0_* and *ΔL* are the initial length of the polymerizing filament and its elongation, respectively, both normalized by the actin sub-unit length. Middle: corresponding kymographs. The sliding *ΔX* and the elongation *ΔL* correspond to the white arrows. Right: Time variation of *ΔX* and *ΔL*. The sliding velocity *v_f_* and the elongation rate *v_p_* are deduced from the slopes of the graphs. Actin fluorescence intensity is represented according to the “Fire” LUT of Image J. Scale bar, 5μm. 1 image/20 sec. **(B)** Myo1b density on the solid substrate or on the supported bilayer deduced from (a) the measurement of the fluorescence intensity of a reference lipid Atto488DOPE at known density in a SLB (supported lipid bilayer), and the comparison of (b) the fluorescence of Myo1b dye Alexa 488 and (c) Atto488DOPE in bulk at known concentrations (see Materials and Methods). The calibration constant *A* is deduced from the slope of a).

**Figure S2:**
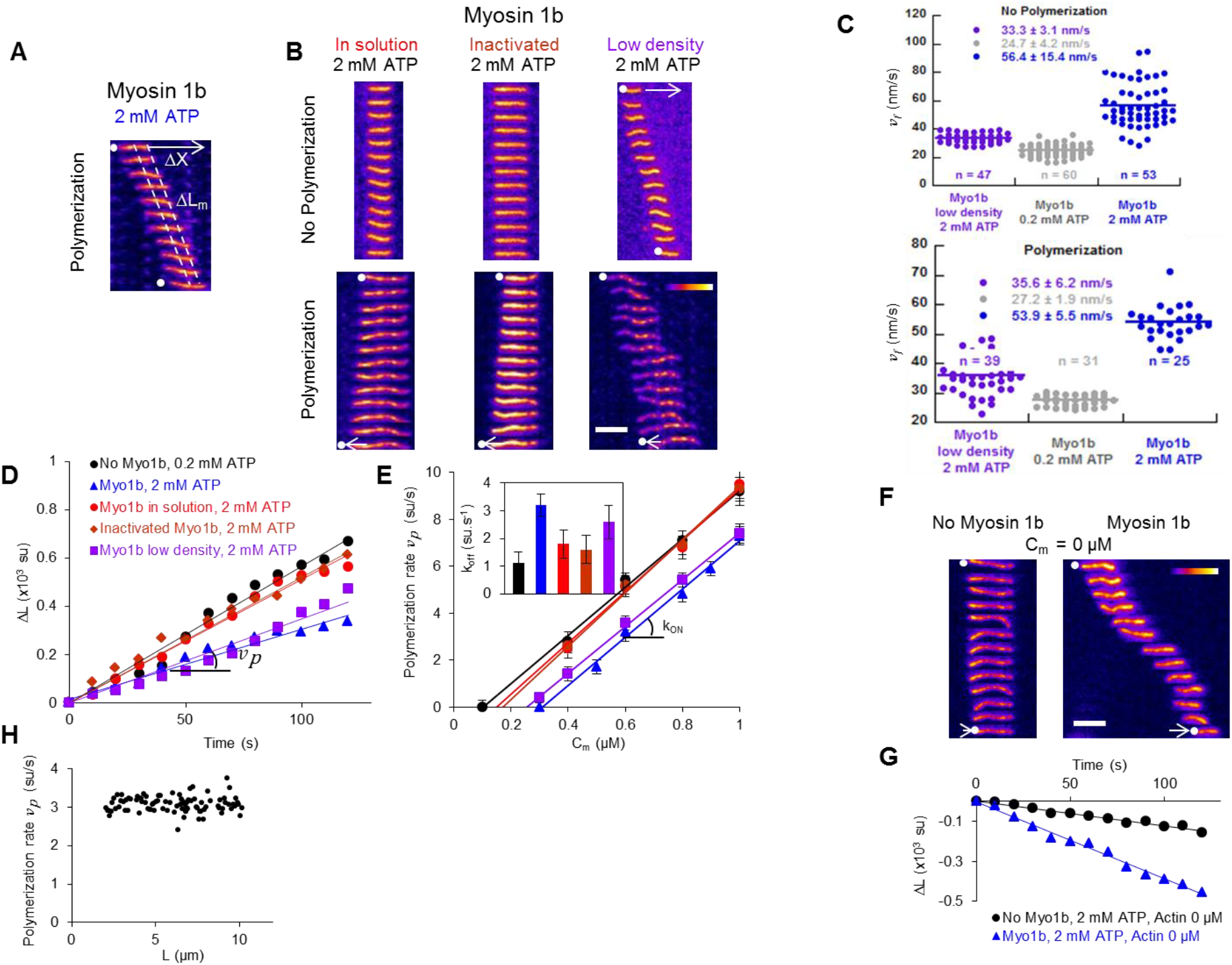
Impact of Myo1b on F-actin pointed end and impact of Myo1b at low density, inactivated, or in solution on the sliding and actin depolymerization at the barbed end. **(A)** Representative kymograph of polymerizing actin filaments, in presence of 0.6 μM G-actin, 2 mM ATP with anchored Myo1b at ≈ 8000 μm^−2^ (see movie S4). The elongation Δ*L_m_* of the filaments at the pointed end (between the 2 dashed white lines) is indicated. Scale bar, 5 μm. 1 image/10 sec. **(B)** Representative kymographs of phalloidin stabilized filaments (top) or polymerizing actin filaments (bottom), in presence of 0.6 μM G-actin, 2 mM ATP, with Myo1b in solution (red), Myo1b without motor activity (brown), with anchored Myo1b at low density (≈500 motor/μm^2^) (purple). (See movies S2, S3 and S5). Scale bar, 5μm. 1image/10 sec. **(C)** Comparison of the distribution of the velocities *v_f_* of stabilized (top) and polymerizing F-actin (bottom) sliding on immobilized high density Myo1b, 2 mM ATP (dark blue) and 0.2 mM ATP (grey) and low density Myo1b, 2 mM ATP (purple). Velocity distributions and average velocities are indicated. Data are represented with a “Dot plot”. The number of analyzed filaments is indicated. **(D)** Δ*L* versus time for the single filaments for the conditions shown in (A) and (B) and in the absence of Myo1b. (E) *v_p_* as a function of G-actin concentration *C_m_* for the different indicated conditions. The fit to the data is the same as in Fig. 2D. Error bars represent s.e.m. (n > 25). Inset: *k_off_* for the different conditions. **(F)** Representative kymographs of depolymerizing actin filaments, in absence of G-actin (*C_m_* = 0 μM), without Myo1b (left) or with anchored Myo1b at ≈8000 motor/μm^2^ (right), with 2 mM ATP. (See movie S7). Scale bar, 5μm. 1 image/10 sec. (G) Δ*L* versus time for the single filaments shown in (F). (H) vp as a function of filament length L for single polymerizing filaments sliding along glass-anchored Myo1b in the presence of 0.6 μM G-actin and 2 mM ATP (n = 90, *v_p_* measured during 30 sec).

**Figure S3:**
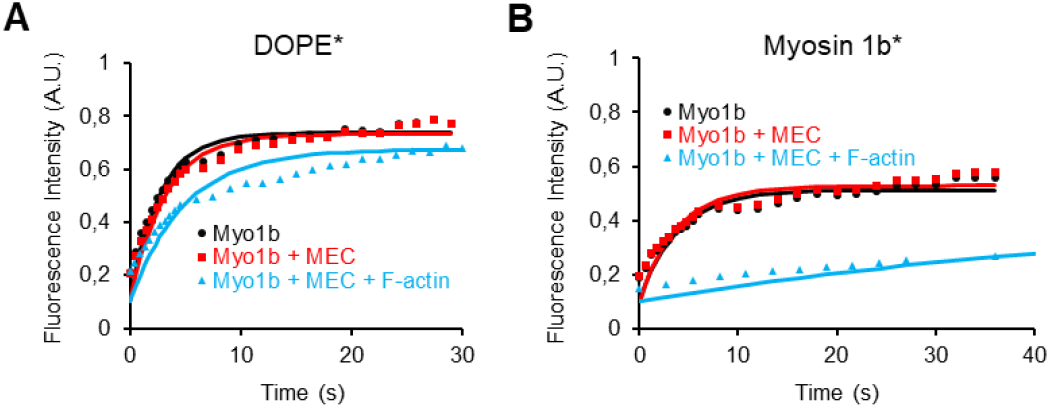
FRAP data of DOPE and Myo1b in a SLB. **(A and B)** Representative FRAP recovery curves (symbols) and the best fit with single exponential (solid line) of Atto488-DOPE (DOPE*) and Alexa488-labelled Myo1b (Myosin 1b*) in a SLB with bound Myo1b, with (in red) or without (in black) 0.3 % methylcellulose (MEC), and in absence or presence (in cyan) of a dense F-actin network.

**Figure S4:**
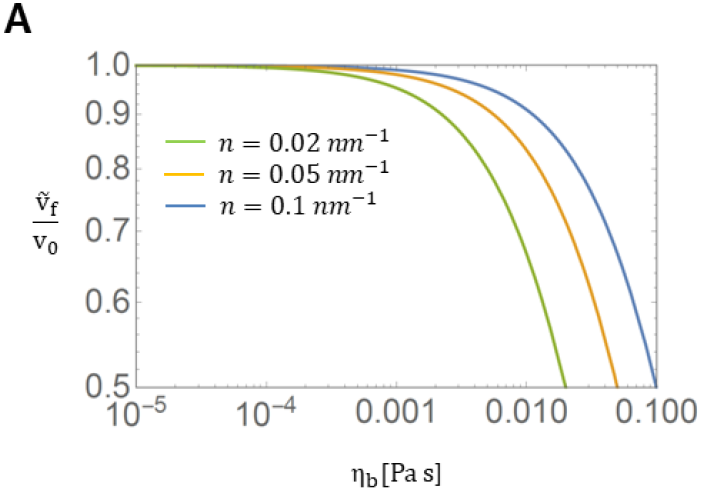
Effect of bulk viscosity on relative velocity of the filament. Velocity of the filament 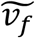, relative to its velocity on a solid substrate *v_0_* (Eq. SE4), for different values of motor density *n*. Increasing the bulk viscosity, relative to the membrane viscosity, induces motion of the motors in the bilayer, hence decreasing the effective velocity of the filament. Increasing the density of molecular motors on the surface increases the effective membrane friction and hence increases the sliding speed.

**Figure S5:**
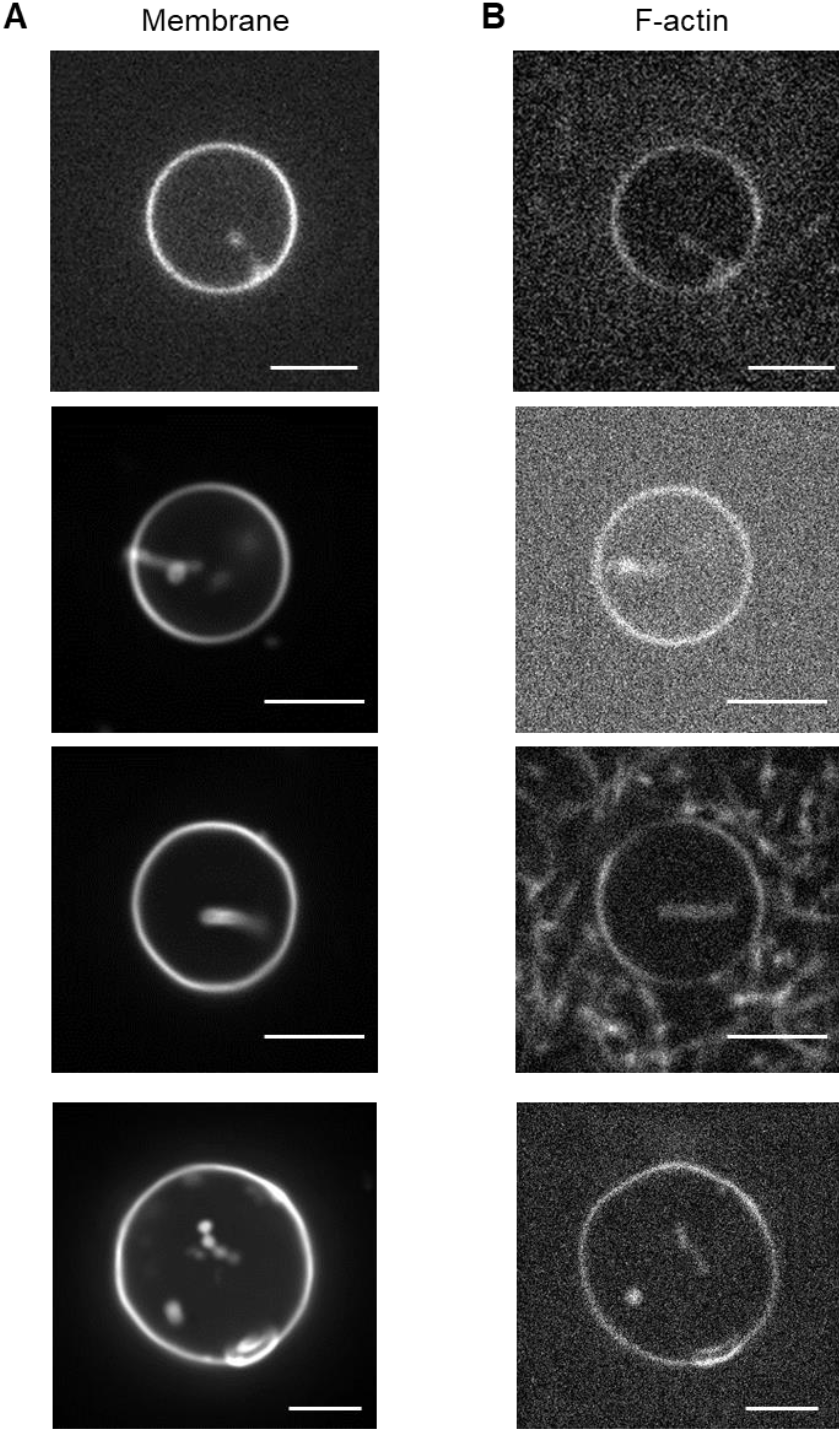
Myo1b bound to giant liposomes produces membrane invagination in presence of stabilized actin filaments. Representative confocal microscopy images of tubules induced by Myo1b bound to a PI(4,5)P_2_-containing GUV in the presence of stabilized actin filaments. We have observed tubulation in the equatorial plane for 16 GUVs over a total of 96. Labeling corresponds to **(A)** 0.3% Texas Red DHPE (mol/mol) and **(B)** stabilized actin filaments with Alexa Fluor 647 phalloidin. Scale bars, 5 μm.

## Legends - Movies

**Movie S1: Stabilized and polymerizing actin filaments sliding along glass-anchored Myo1b (corresponding to Fig. S1A).**

Sliding of stabilized filaments labeled with phalloidin-Alexa547 (a) and polymerizing filaments (b) with 1.2 μM actin in bulk (10 % Alexa594 labeled), along Myo1b (8000 μm^−2^) at 2 mM ATP. White arrow indicates the initial position of the filament corresponding to the kymograph in Fig. S1A. Note that no filament breaking is observed during these experiments. Scale bar 5 μm. Time in s.

**Movie S2: Effect of the ATP concentration and Myo1b density on stabilized actin filaments sliding along glass-anchored Myo1b (corresponding to Figs. 2A and S2B).**

Filaments stabilized with phalloidin-Alexa547 as a function of ATP concentration (0.2 and 2 mM ATP), without Myo1b (a), sliding on Myo1b at high density (8000 μm^−2^) (b, c) or at low density (400 μm^−2^) (d). Wide field movies followed by single filament movies corresponding to kymographs shown in Fig. 2A-Top panel and Fig. S2B-Top panel. White arrows indicate the initial position of single filaments. Scale bar 5 μm. Time in s.

**Movie S3: Effect of ATP concentration and Myo1b density on polymerizing actin filaments sliding along glass-anchored Myo1b (corresponding to Fig. S2A).**

Polymerizing filaments as a function of ATP concentration as a function of ATP concentration (0.2 and 2 mM ATP), without Myo1b (a), sliding on Myo1b at high density (8000 μm^−2^) (b, c) or at low density (400 μm^−2^) (d). Wide field movies followed by single filament movies corresponding to kymographs shown in Fig. 2A-Bottom panel and Fig. S2B-Bottom panel. White arrows indicate the initial position of single filaments. Scale bar 5 μm. Time in s.

**Movie S4: No effect at pointed-end on polymerizing actin filaments sliding along glass-anchored Myo1b.**

Polymerizing actin filaments with 0.6 μM actin (10 % Alexa594 labeled), along Myo1b (8000 μm^−2^) at 2 mM ATP. The white arrow indicates the initial position of the filament shown in Fig. S2A. Scale bar 5 μm. Time in s.

**Movie S5: Stabilized and polymerizing actin filaments with Myo1b in bulk or inactivated (corresponding to Fig. S2B).**

Filaments stabilized with phalloidin-Alexa547 and polymerizing actin filament, with 0.6 μM actin (10 % Alexa594 labeled), 2 mM ATP, 300 nM Myo1b in the bulk, or bound but inactivated. Note that in the bottom left movie, one filament seems to move but it suddenly appears in the field of view while sedimenting. Frames correspond to the kymographs shown in Fig. S2B; arrows indicate the initial position of these filaments. Scale bar 5 μm. Time in s.

**Movie S6: Stabilized and polymerizing actin filaments sliding along glass-anchored MyoII (corresponding to Fig. 2A).**

Filaments stabilized with phalloidin-Alexa547 or polymerizing at 0.6 μM actin (10 % Alexa594 labeled), sliding along MyoII at 2 mM ATP. Frames correspond to the kymographs shown in Fig. 2A; arrows indicate the initial position of these filaments. Scale bar 5 μm. Time in s.

**Movie S7: Impact of Myo1b on the depolymerization of actin filaments in the absence of G-actin in the bulk (corresponding to Fig. S2F).**

Filaments depolymerizing in the absence of G-actin in bulk, without Myo1b, or sliding along Myo1b, at 2 mM ATP. Frames correspond to the kymographs shown in Fig. S2F; arrows indicate the initial position of these filaments. Scale bar 5 μm. Time in s.

**Movie S8: Stabilized and polymerizing actin filaments sliding along Myo1b bound to SLBs (corresponding to Fig. 3B).**

Sliding of filaments stabilized with phalloidin-Alexa547 or polymerizing at 0.6 μM actin (10 % Alexa594 labeled), along Myo1b bound to SLBs (≈ 8500 μm^−2^) at 2 mM ATP. Wide field movies followed by single filament movies corresponding to kymographs shown in Fig. 3B. White arrows indicate the initial position of single filaments. Scale bar 5 μm. Time in s.

## References

1. Köster, D. V. & Mayor, S. Cortical actin and the plasma membrane: inextricably intertwined. Curr. Opin. Cell Biol. 38, 81–89 (2016).

2. Pyrpassopoulos, S., et al. Force Generation by Membrane-Associated Myosin-I. Sci. Rep. 6, 25524 (2016).

3. McIntosh, B. B. & Ostap, E. M. Myosin-I molecular motors at a glance. J. Cell Sci. 129, 2689–2695 (2016).

4. Almeida, C. G., et al. Myosin 1b promotes the formation of post-Golgi carriers by regulating actin assembly and membrane remodelling at the trans-Golgi network. Nat. Cell Biol. 13, 779–789 (2011).

5. Gupta, P., et al. Myosin 1E localizes to actin polymerization sites in lamellipodia, affecting actin dynamics and adhesion formation. Biol. Open 2, 1288–1299 (2013).

6. Iuliano, O., et al. Myosin 1b promotes axon formation by regulating actin wave propagation and growth cone dynamics. J. Cell Biol. 217, 2033 (2018).

7. Joensuu, M., et al. ER sheet persistence is coupled to myosin 1c–regulated dynamic actin filament arrays. Mol. Biol. Cell 25, 1111–1126 (2014).

8. Laakso, J. M., Lewis, J. H., Shuman, H. & Ostap, E. M. Myosin I can act as a molecular force sensor. Science 321, 133–136 (2008).

9. Greenberg, M. J., Lin, T., Goldman, Y. E., Shuman, H. & Ostap, E. M. Myosin IC generates power over a range of loads via a new tension-sensing mechanism. Proc. Natl Acad. Sci. USA 109, E2433–E2440 (2012).

10. Lewis, J. H., Lin, T., Hokanson, D. E. & Ostap, E. M. Temperature dependence of nucleotide association and kinetic characterization of Myo1b. Biochemistry 45, 11589–11597 (2006).

11. Carlier, M.-F., Pernier, J., Montaville, P., Shekhar, S. & Kühn, S. Control of polarized assembly of actin filaments in cell motility. Cell. Mol. Life Sci. 72, 3051–3067 (2015).

12. Veigel, C., Molloy, J. E., Schmitz, S. & Kendrick-Jones, J. Load-dependent kinetics of force production by smooth muscle myosin measured with optical tweezers. Nat. Cell Biol. 5, 980 (2003).

13. Komaba, S. & Coluccio, L. M. Localization of Myosin 1b to Actin Protrusions Requires Phosphoinositide Binding. J. Biol. Chem. 285, 27686–27693 (2010).

14. Guo, L., et al. Molecular diffusion measurement in lipid bilayers over wide concentration ranges: a comparative study. Chem Phys Chem 9, 721–728 (2008).

15. Pyrpassopoulos, S., Feeser, E. A., Mazerik, J., Tyska, M. & Ostap, M. Membrane-bound Myo1c powers asymmetric motility of actin filaments. Curr. Biol. 22, 1688–1692 (2012).

16. Grover, R., et al. Transport efficiency of membrane-anchored kinesin-1 motors depends on motor density and diffusivity. Proc. Natl Acad. Sci. USA 113, E7185 (2016).

17. Vogel, S. K., Petrasek, Z., Heinemann, F. & Schwille, P. Myosin motors fragment and compact membrane-bound actin filaments. eLife 2, e00116 (2013).

18. Murrell, M. & Gardel, M. L. F-actin buckling coordinates contractility and severing in a biomimetic actomyosin cortex Proc. Natl Acad. Sci. USA 109, 20820–20825 (2012).

19. Kovar, D. R., Kuhn, J. R., Tichy, A. L. & Pollard, T. D. The fission yeast cytokinesis formin Cdc12p is a barbed end actin filament capping protein gated by profilin. J. Cell Biol. 161, 875 (2003).

20. Romero, S., et al. Formin Is a Processive Motor that Requires Profilin to Accelerate Actin Assembly and Associated ATP Hydrolysis. Cell 119, 419–429 (2004).

21. Johnston, A. B., Collins, A. & Goode, B. L. High-speed depolymerization at actin filament ends jointly catalysed by Twinfilin and Srv2/CAP. Nat. Cell Biol. 17, 1504 (2015).

22. Wioland, H., et al. ADF/Cofilin accelerates actin dynamics by severing filaments and promoting their depolymerization at both ends. Curr. Biol. 27, 1956–1967.e1957 (2017).

23. Varga, V., Leduc, C., Bormuth, V., Diez, S. & Howard, J. Kinesin-8 Motors Act Cooperatively to Mediate Length-Dependent Microtubule Depolymerization. Cell 138, 1174–1183 (2009).

24. Moores, C. A. & Milligan, R. A. Lucky 13-microtubule depolymerisation by kinesin-13 motors. J. Cell Sci. 119, 3905–3913 (2006).

25. Crevenna, A. H., et al. Side-binding proteins modulate actin filament dynamics. eLife 4, e04599 (2015).

26. Greenberg, M. J., Lin, T., Shuman, H. & Ostap, E. M. Mechanochemical tuning of myosin-I by the N-terminal region. Proc. Natl Acad. Sci. USA 112, E3337–E3344 (2015).

27. Ennomani, H., et al. Architecture and Connectivity Govern Actin Network Contractility. Curr. Biol. 26, 616–626 (2016).

28. Freeman, S. A., et al. Toll-like receptor ligands sensitize B-cell receptor signalling by reducing actin-dependent spatial confinement of the receptor. Nat. Commun. 6, 6168 (2015).

29. Dürre, K., et al. Capping protein-controlled actin polymerization shapes lipid membranes. Nat. Commun. 9, 1630 (2018).

30. Noguchi, T., Lenartowska, M. & Miller, K. G. Myosin VI stabilizes an actin network during Drosophila spermatid individualization. Mol. Biol. Cell 17, 2559–2571 (2006).

31. Loubéry, S., Delevoye, C., Louvard, D., Raposo, G. & Coudrier, E. Myosin VI Regulates Actin Dynamics and Melanosome Biogenesis. Traffic 13, 665–680 (2012).

32. Ciobanasu, C., Faivre, B. & Le Clainche, C. Actomyosin-dependent formation of the mechanosensitive talin–vinculin complex reinforces actin anchoring. Nat. Commun. 5, 3095 (2014).

33. Pollard, T. D. Myosin Purification and Characterization. In: Methods in Cell Biology (ed^(eds Wilson L). Elsevier Inc. (1982).

34. Yamada, A., et al. Catch-bond behaviour facilitates membrane tubulation by non-processive myosin 1b. Nat. Commun. 5, 3624 (2014).

35. Weinberger, A., et al. Gel-assisted formation of Giant Unilamellar Vesicles. Biophys. J. 105, 154–164 (2013).

36. Galush, W. J., Nye, J. A. & Groves, J. T. Quantitative fluorescence microscopy using supported lipid bilayer standards. Biophys. J. 95, 2512–2519 (2008).

37. Sorre, B., et al. Nature of curvature-coupling of amphiphysin with membranes depends on its bound density. Proc. Natl Acad. Sci. USA 109, 173–178 (2012).

38. Soumpasis, D. M. Theoretical analysis of fluorescence photobleaching recovery experiments. Biophys. J. 41, 95–97 (1983).

39. Vilfan, A., Frey, E. & Schwabl, F. Force-velocity relations of a two-state crossbridge model for molecular motors. Europhys. Lett. 45, 283–289 (1999).

40. Saffman, P. G. & Delbrück, M. Brownian motion in biological membranes. Proc. Natl Acad. Sci. USA 72, 3111–3113 (1975).

